## Supplementary Materials for "Promoter-anchored chromatin interactions predicted from genetic analysis of epigenomic data"

Wu *et al.*

**Contents**

**Figure S1 to S12**
**Supplementary Note 1-3**
**References**

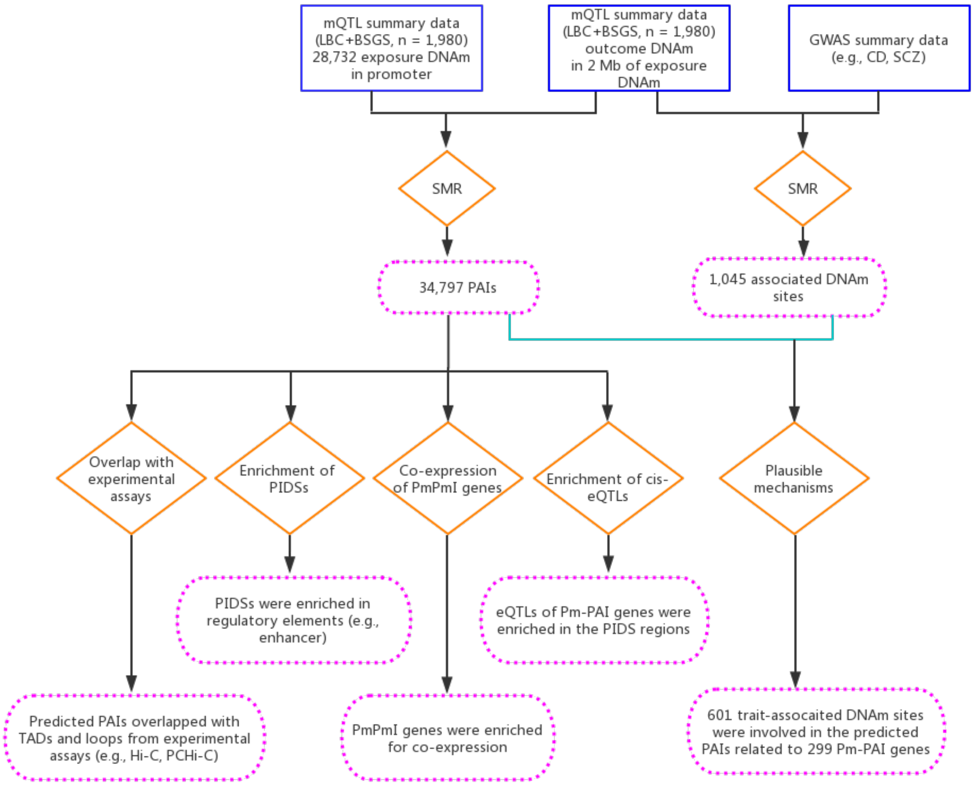


**Figure S1** Schematic overview of this study.

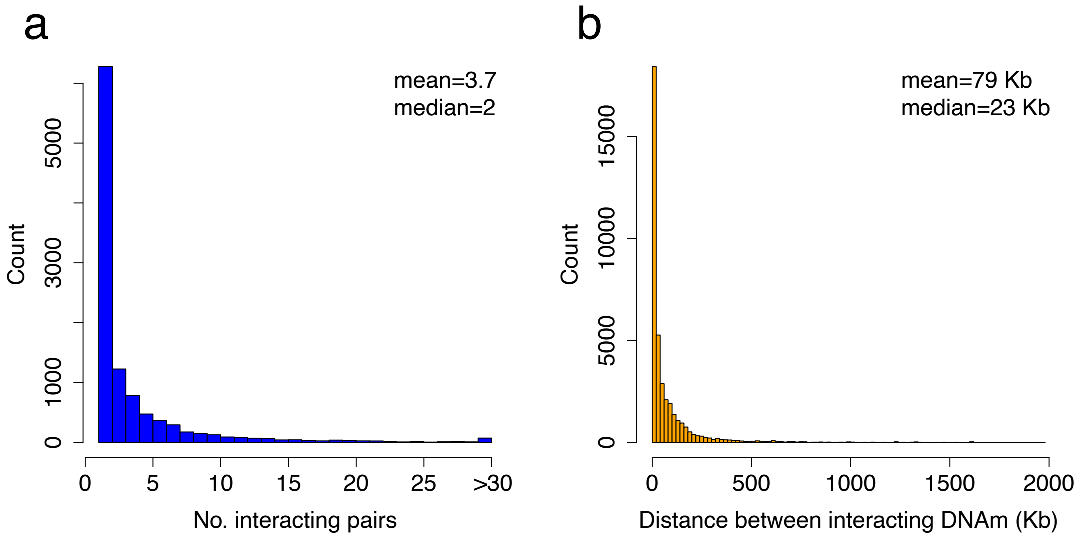


**Figure S2** Summary of the predicted PAIs. Panel a): distribution of the number of PIDSs (promoter interacting DNAm sites) for each bait probe (located in the promoter of a gene). Panel b): distribution of physical distances between pairwise interacting DNAm sites of the significant PAIs.

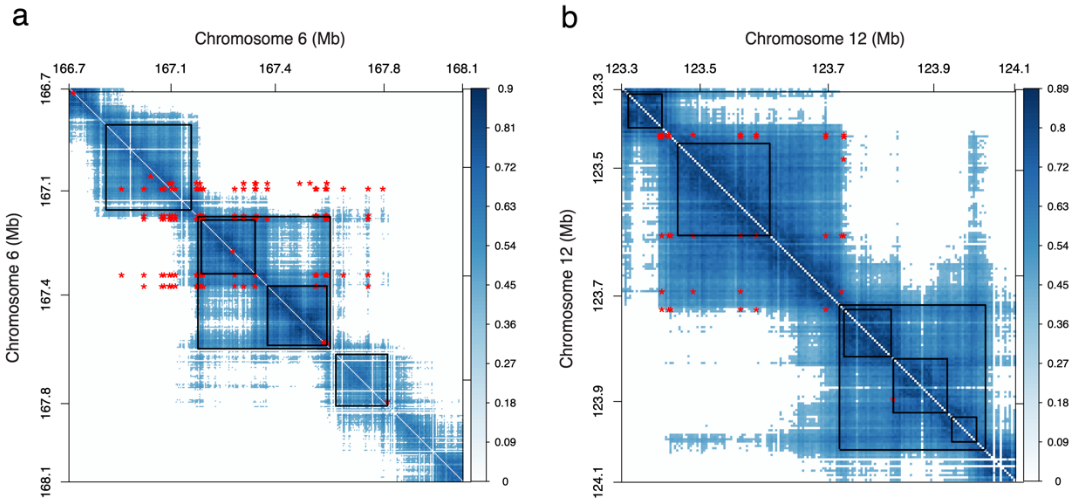


**Figure S3** Overlap of the predicted PAIs with TADs annotated from the Rao *et al.*[1] Hi-C data. Panel a): a heatmap of the predicted PAIs (red asterisks) and chromatin interactions with correlation scores > 0.4 (blue dots) identified by Hi-C from Grubert *et al.*[2] in a 1.38 Mb region on chromosome 6. Black squares are the TADs identified by Rao *et al.*[1]. Only 41.5% of the predicted PAIs in this region showed overlap with the TADs. This region harbours the *RPS6KA2* locus as shown in **Fig. 6**. Panel b): a heatmap of the predicted PAIs (red asterisks) and chromatin interactions with correlation scores > 0.4 (blue dots) identified by Hi-C from Grubert *et al.*[2] in a 0.81 Mb region on chromosome 12. Black squares are the TADs identified by Rao *et al.*[1]. The predicted PAIs were highly consistent with the chromatin interactions identified by Hi-C. This region harbours the *ABCB9* locus as shown in **Fig. S7**. The heatmap is asymmetric for the PAIs with the *x*- and *y*-axes representing the physical positions of “outcome” and “exposure” probes respectively.

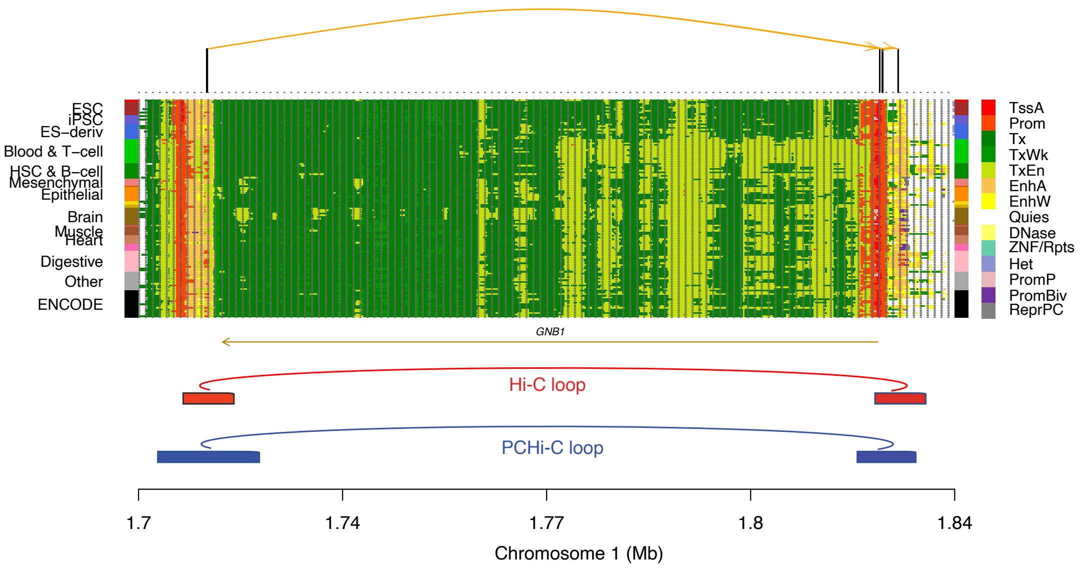


**Figure S4** Validation of the predicted PAIs by the enhancer-promoter interactions identified by previous experimental assays at the *GNB1* locus. Shown are the 14 REMC chromatin state annotations with the PAIs labelled on the top and the chromatin loops labelled on the bottom. The annotated Hi-C (red) and PCHi-C (blue) loops are identified by Rao *et al.*[1] and Jung *et al.*[3], respectively, in GM12878 cell lines.

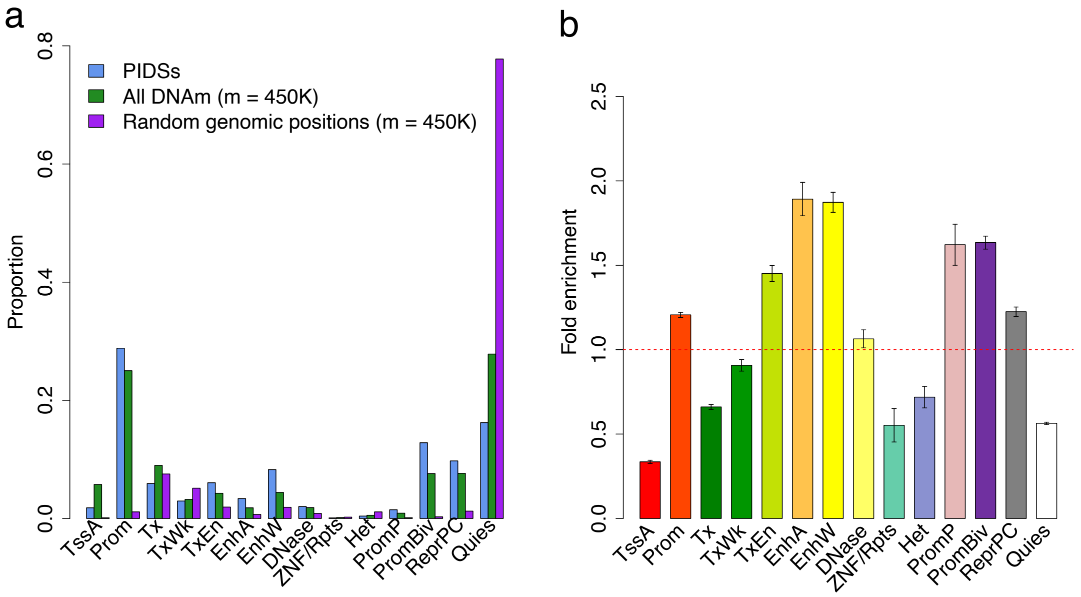


**Figure S5** Distribution of DNAm sites in the 14 REMC functional annotation categories. Panel a): distribution of the PIDSs identified in this study (blue), all DNAm probes on an Illumina 450K methylation array (green), and 450K genomic positions randomly sampled across the genome (purple) in the 14 annotation categories. Panel b): enrichment of the PIDSs in the 14 annotation categories when the DNAm pairs within a promoter were included in the PAI analysis. Fold enrichment: a ratio of the proportion of PIDSs in an annotation category to the mean of the control sets. In the analysis presented in Figure 4b, DNAm pairs within a promoter were excluded from the SMR analysis, which was likely the cause of the depletion of the PIDSs in promoters. To test this hypothesis, we added the within-promoter DNAm pairs back in analysis. Each error bar represents the standard deviation of an estimate obtained from 1,000 control sets.

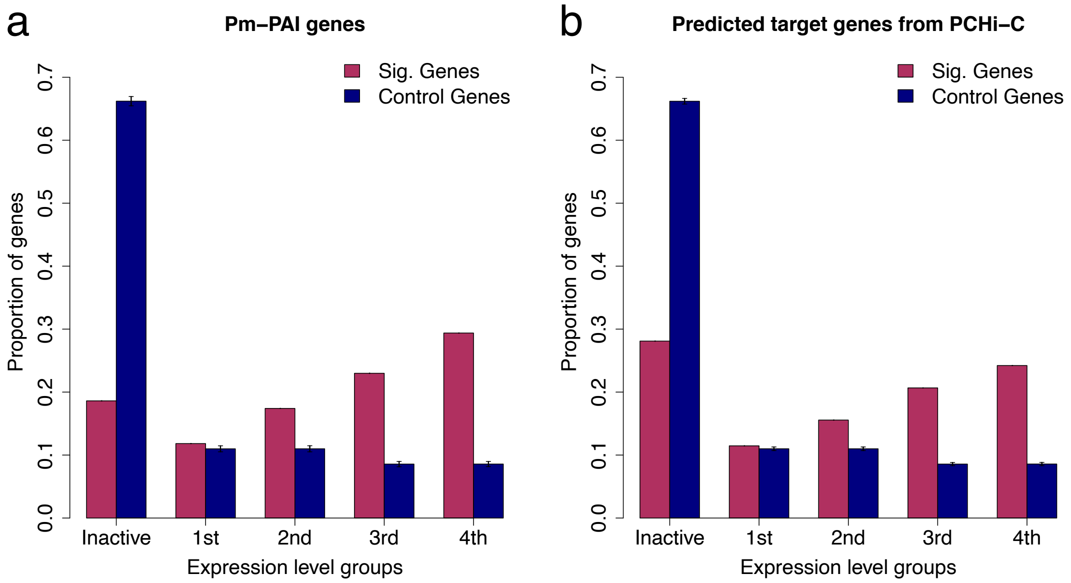


**Figure S6** Proportion of the predicted genes involved in chromatin interactions in five gene activity groups. Panel a): proportion of the Pm-PAI genes in five gene activity groups. Panel b): proportion of the predicted target genes whose promoters were involved in the physical interactions identified by PCHi-C from the Jung *et al.* study[3] in five gene activity groups. In the analysis presented in Figure 4d, the control genes were randomly sampled from the genes whose promoters were tested in the PAI analysis. However, this enrichment analysis is conservative because the ascertainment of genes with promoter DNAm sites tested in SMR would potentially lead to upward biases in expression levels of the control genes. Therefore, we performed an additional enrichment analysis by randomly resampling the control genes from all the genes available in the GTEx blood samples. The genes were divided into five gene activity groups with the first group being the inactive group (TPM < 0.1) together with four quartiles defined based on the expression levels of all genes in the GTEx blood samples. Each error bar represents the standard deviation estimated from the control genes in 1,000 random samples.

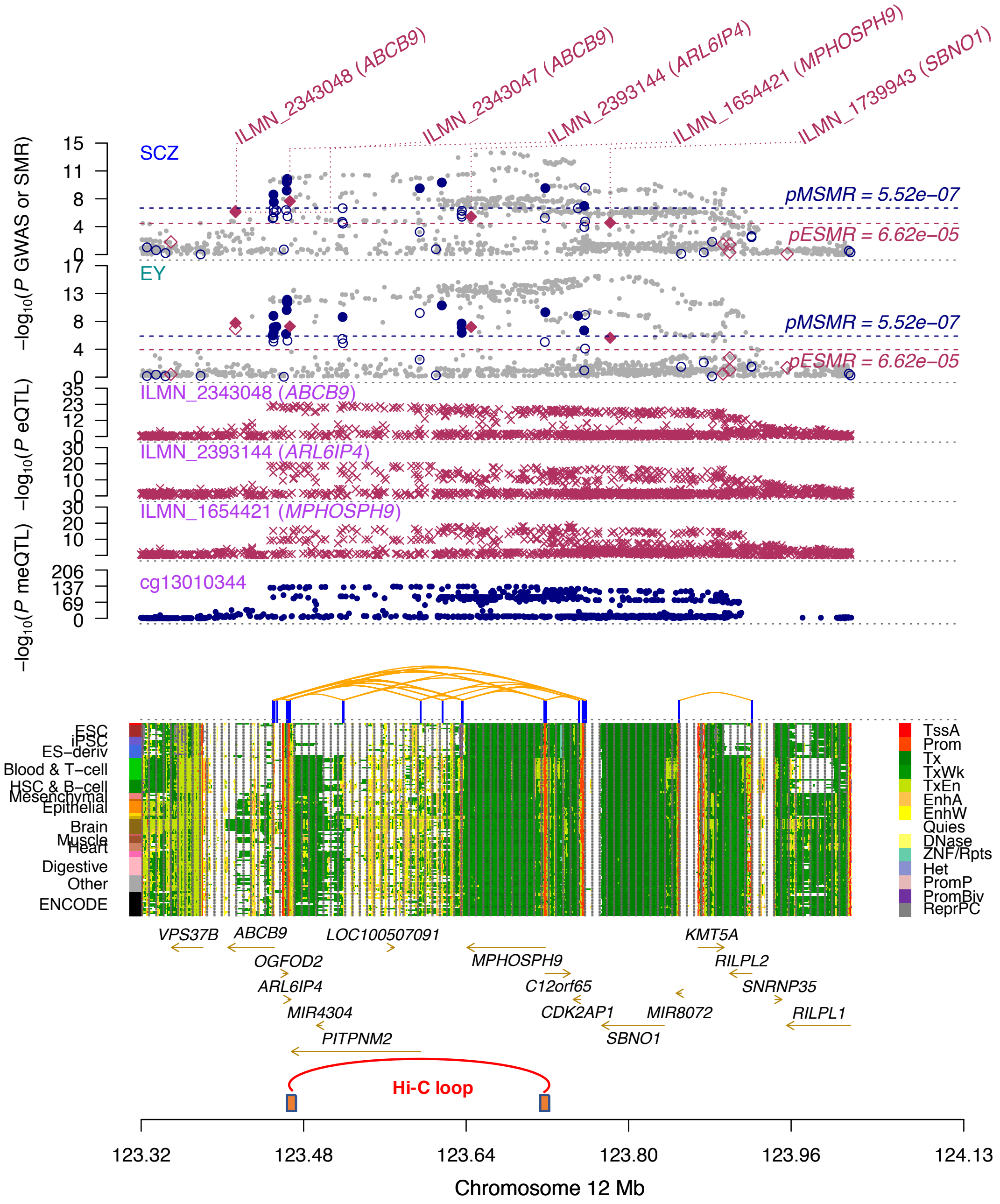


**Figure S7** A shared PIDS region with eQTLs predicted to interact with the promoters of multiple genes (i.e., *ABCB9*, *ARL6IP4*, *MPHOSPH9*). The top two plots show -$\log_{10}$(*P* values) of SNPs from the GWAS meta-analyses (grey dots) for schizophrenia (SCZ) and educational years (EY). Red diamonds and blue circles represent -$\log_{10}$(*P* values) from SMR tests for associations of gene expression and DNAm with SCZ and EY, respectively. Solid diamonds and circles are the probes not rejected by the HEIDI test. The following three plots show -$\log_{10}$(*P* values) of SNP associations for the gene expression probes ILMN_2343048 (tagging *ABCB9*), ILMN_2393144 (tagging *ARL6IP4*), and ILMN_1654421 (tagging *MPHOSPH9*) from the CAGE study. The sixth plot shows -$\log_{10}$(*P* values) of SNP associations for the DNAm probe cg13010344 from the mQTL meta-analysis. The heatmap-like panel on the bottom shows the 14 REMC annotations with the significant PAIs annotated by orange curved lines on the top (see **Fig. S3b** for the overlap of the predicted PAIs with Hi-C data) and the Hi-C loop identified by Rao *et al.*[1] annotated on the x-axis (two orange bars connected by a red curved line).

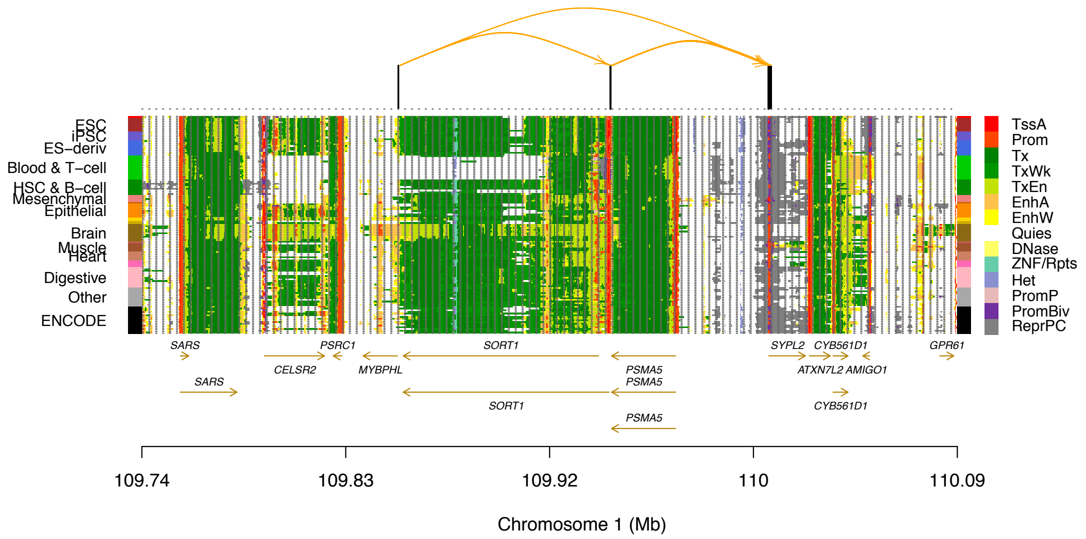


**Figure S8** Predicted PAIs at the *SORT1* locus. Shown are the 14 REMC chromatin state annotations with the significant PAIs labelled on the top.

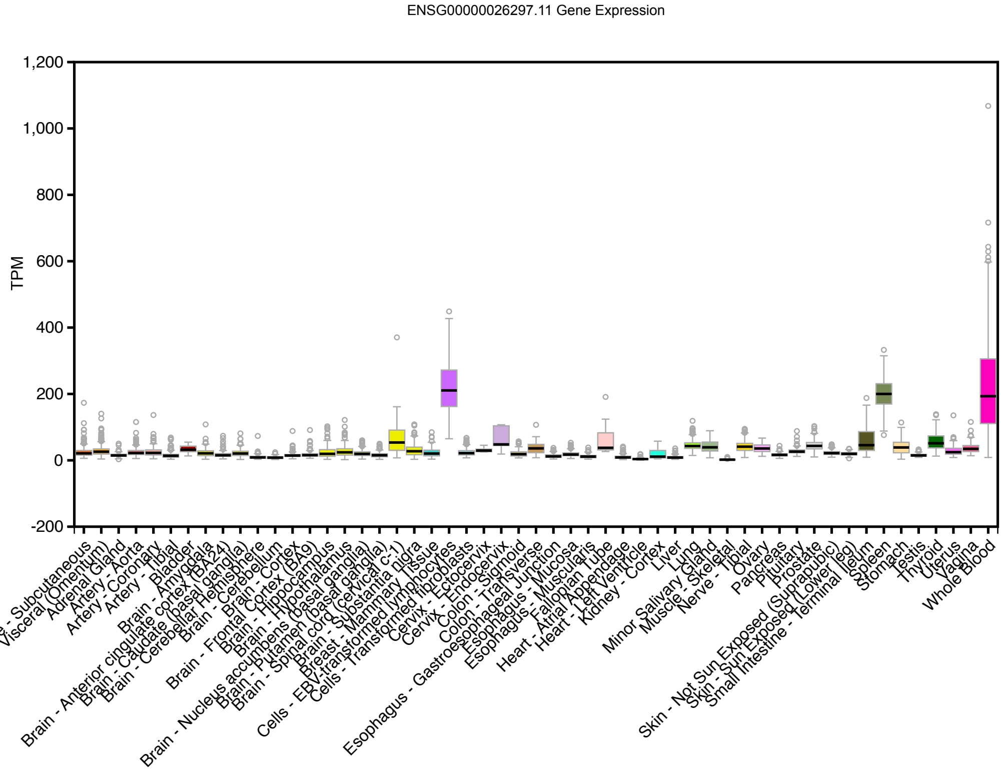


**Figure S9** Expression (measured by TPM) of *RNASET2* (ENSG00000026297.11) in 53 tissues of the GTEx (V7 data).

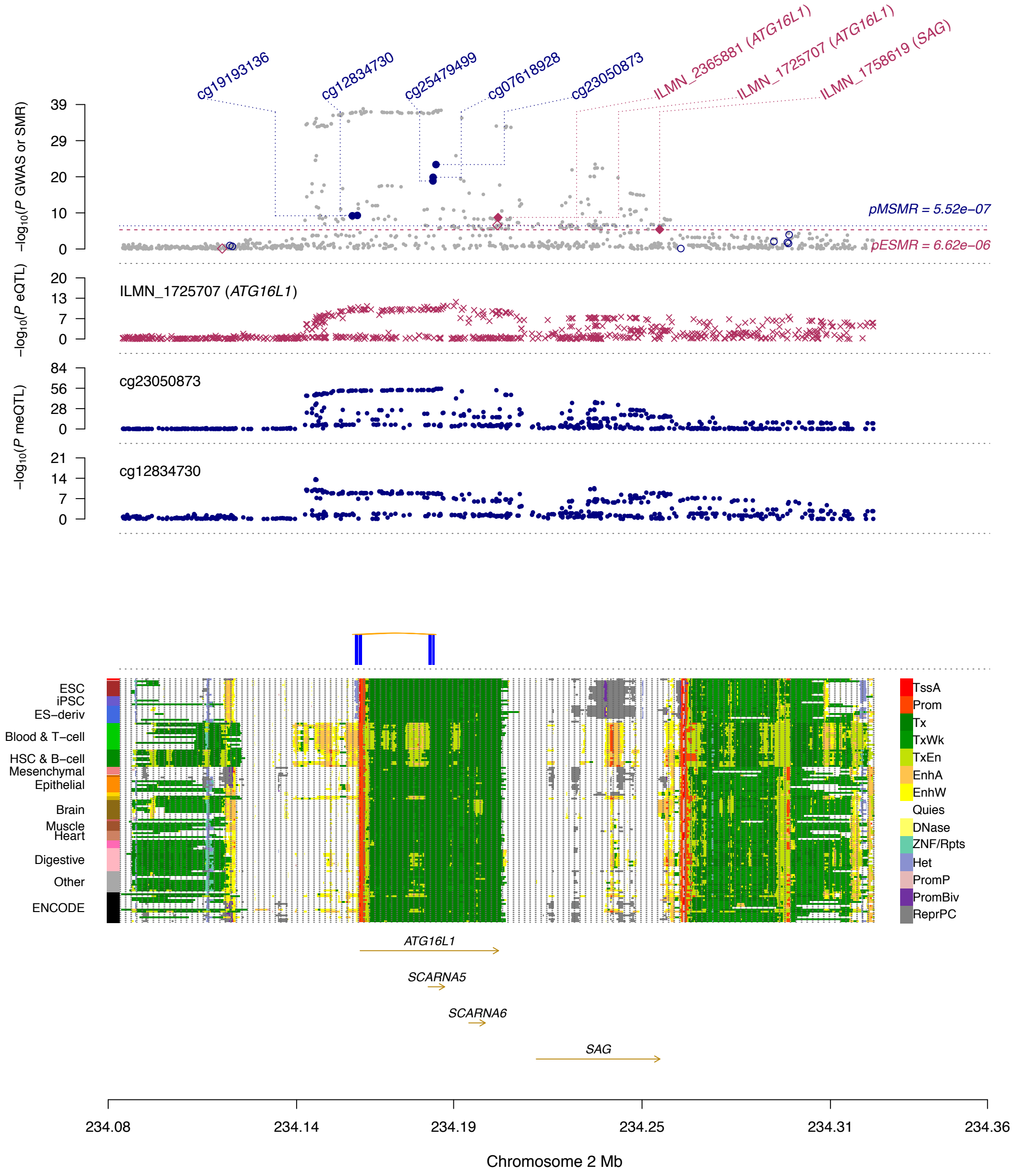


**Figure S10** Prioritizing gene and functional regions at the *ATG16L1* locus for Crohn's disease (CD). The top plot shows -$\log_{10}$($P$ values) of SNPs from the GWAS meta-analysis (grey dots) for CD[4]. Red diamonds and blue circles represent -$\log_{10}$($P$ values) from the SMR tests for associations of gene expression and DNAm with CD respectively. Solid diamonds are the probes not rejected by the HEIDI test. The second plot shows -$\log_{10}$($P$ values) of SNP associations for the gene expression probe ILMN_1725707 (tagging *ATG16L1*) from the CAGE study. The third plot shows -$\log_{10}$($P$ values) of SNP associations for the DNAm probes cg23050873 and cg12834730. The plot in the middle shows the significant PAIs between pairwise DNAm sites. The bottom plot shows the 14 REMC chromatin state annotations.

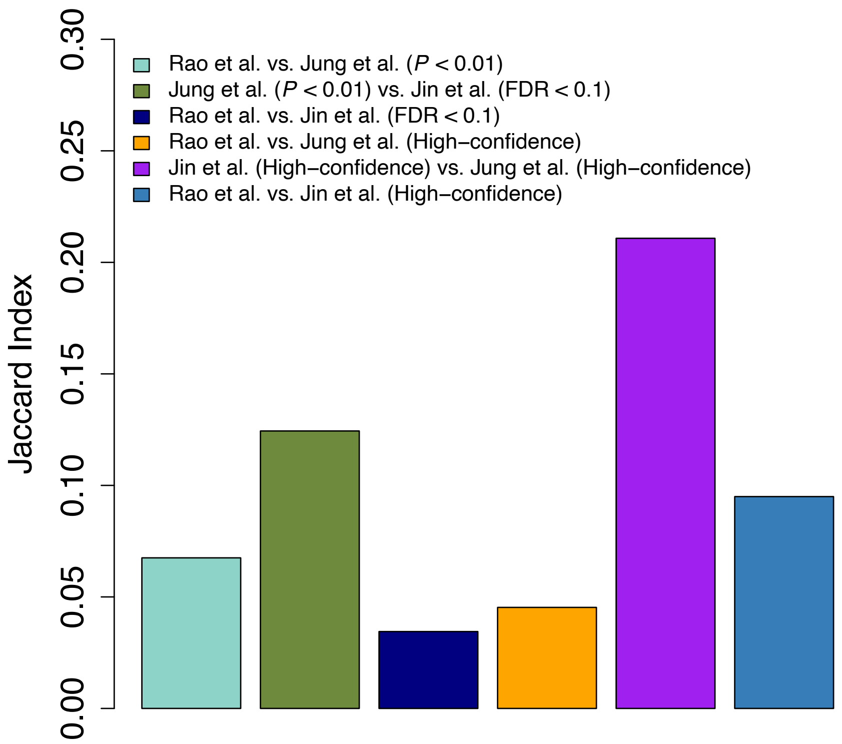


**Figure S11** Concordances of the identified chromatin loops between pairwise Hi-C datasets. The chromatin interactions from Jung *et al.*[3] are defined at $P < 0.01$ and those from Jin *et al.*[5] are defined at a false discovery rate (FDR) < 0.01. The high-confidence Hi-C loops from Jung *et al.* and Jin *et al.* are defined as the top *m* chromatin loops with the smallest Hi-C *P* values, where *m* is the number of Hi-C loops identified in Rao *et al.*[1] ($m$ = 9,448). The Jaccard Index of each pair of datasets is defined as the ratio of the size of the intersection to that of the union.

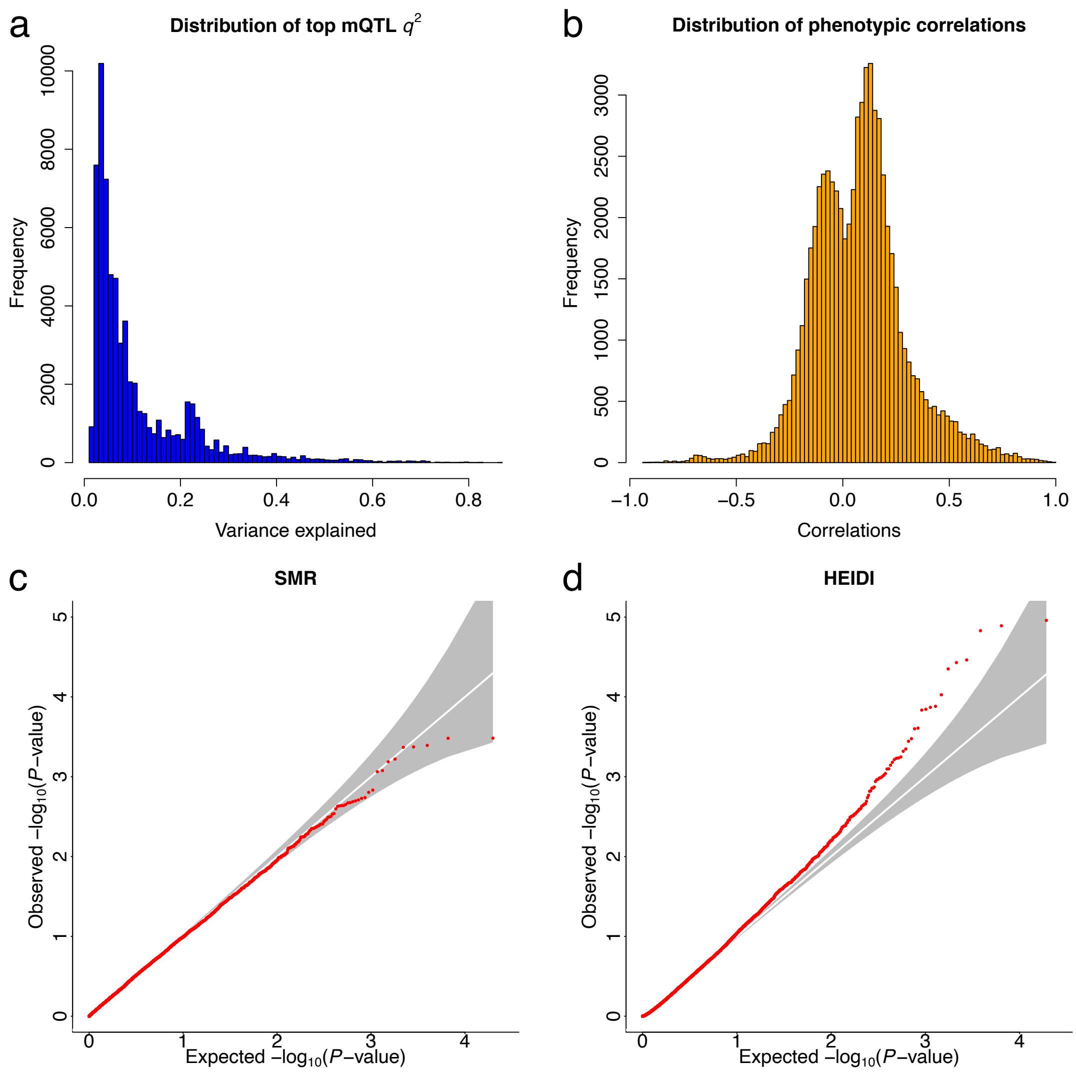


**Figure S12** Calibrating the test-statistics from SMR and HEIDI by simulation in overlapping samples. Details of the simulation can be found in **Supplementary Note 1**. Panel a): distribution of the variance in DNAm explained by the top-associated mQTL for each exposure probe. Panel b): distribution of correlations of DNAm levels between pairwise CpG sites (included in the PAI analysis) computed from LBC data[6] (**Methods**). Panel c): QQ plot of *P* values from the SMR test under the null model (i.e., there is no association between the two DNAm sites). Panel d): QQ plot of *P* values from the HEIDI test under the pleiotropic model (i.e., the two DNAm sites are associated due to the same causal variant).

**Supplementary Note 1. The reasons why the predicted PAIs are relatively sparse**

There are several reasons why the predicted PAIs are relatively sparse. First, although the Illumina 450K methylation array has a genome-wide coverage, the probes cover only a limited proportion of the regulatory elements. Second, SMR requires the exposure probe with at least an mQTL at $P_{mQTL}$ < 5e-8, and we limited the exposure probes in promoter regions, resulting in only a small proportion of exposure DNAm probes (m=28,732, ~6.5%) being included in the SMR & HEIDI analysis. Third, to control for false positives, we applied an experiment-wise SMR significance threshold (i.e., $P_{\mathrm{SMR}}$ < 1.76e-9) to correct for multiple testing and a stringent HEIDI threshold (i.e., $P_{\mathrm{HEIDI}}$ < 0.01) to reject SMR associations due to linkage. However, despite the relatively sparse distribution of the predicted PAIs across the genome, the number of predicted PAIs (*m*=34,797) is comparable to the loops identified by experimental assays such as Hi-C and PCHi-C. For example, there are only ~10,000 Hi-C loops identified from Rao *et al.*[1] and ~80,000 PCHi-C loops identified from Jung *et al.*[3].

**Supplementary Note 2. Simulations in overlapping samples**

The SMR and HEIDI tests assume that the estimate of SNP effect on exposure is independent of that on outcome. This assumption can be violated if the sample from which the SNP effect on exposure was estimated overlapped with that from which the SNP effect on outcome was estimated, and there is phenotypic correlation between exposure and outcome in the overlapping samples[7]. To investigate whether the SMR and HEIDI test-statistics were biased by sample overlap, we performed simulations under the null and pleiotropic models based on the whole-genome sequencing (WGS) data from the UK10K project[8]. We included only unrelated individuals (*n* = 3,642) and ~8.3 million SNPs with minor allele frequency (MAF) > 0.01 and Hardy-Weinberg Equilibrium (HWE) *P* value > $1\times10^{-6}$.

**Simulation under the null model**

We first simulated DNAm levels at two probes in the same sample (i.e., complete sample overlap) under the null model (i.e., there is no association between the DNAm levels of the two probes) and investigated whether the SMR statistics were biased or not. To do so, we randomly sampled SNPs within a 1 Mb region of the genome and chose one SNP at random as the causal SNP from the sampled SNPs. We then simulated the DNAm levels of 3,642 individuals at one probe ($\boldsymbol{m_1}$) based on model $\boldsymbol{m_1} = \boldsymbol{z}b_{zm1} + \boldsymbol{e}_{m1}$, where $\boldsymbol{z}$ is a vector of genotype of the causal SNP, $b_{zm1}$ is the effect of the causal SNP on DNAm level $b_{zm1} \sim N(0, R_{zm1}^2)$ with $R_{zm1}^2$ being the proportion of variance in $\boldsymbol{m_1}$ explained by the causal SNP, and $\boldsymbol{e}_{m1} \sim N(0, \sigma_{e_{m1}}^2)$ with $\sigma_{e_{m1}}^2 = var(\boldsymbol{Z}b_{zm1})(1/R_{zm1}^2 - 1)$. To generate data under the null model, we generated the DNAm levels of the other probe ($\boldsymbol{m_2}$) based on model $\boldsymbol{m}_2 = \boldsymbol{e}_{m2}$ , where $\boldsymbol{e}_{m2} \sim N(0, \sigma_{e_{m2}}^2)$ with $\sigma_{e_{m2}}^2 = 1$ . Correlation of errors in estimating the SNP effects ($r_{\mathbf{e}}$) may occur due to sample overlap ($\rho$) and phenotypic correlation ($r_p$). To mimic this, we generated residuals ($\boldsymbol{e}$) of the two probes from a multivariate normal distribution, $\boldsymbol{e} = \begin{pmatrix} \boldsymbol{e}_{m1} \\ \boldsymbol{e}_{m2} \end{pmatrix} \sim N\left(\begin{pmatrix} 0 \\ 0 \end{pmatrix}, \begin{pmatrix} \sigma_{e_{m1}}^2 & r_p\sigma_{e_{m1}}\sigma_{e_{m2}} \\ r_p\sigma_{e_{m1}}\sigma_{e_{m2}} & \sigma_{e_{m2}}^2 \end{pmatrix}\right)$, where $r_p$ and $R_{zm1}^2$ were randomly sampled from the observed distributions computed from data used in the PAI analysis in the LBC cohorts. We then performed a regression analysis to detect the top associated mQTL for each simulated probe, and ran an SMR analysis for each pair of simulated probes. We repeated the simulation 5,000 times to quantify the inflation/deflation of SMR test-statistics under this simulation scenario (**Fig. S12c**).

**Simulation under a pleiotropic model**

To examine the distribution of the HEIDI test-statistics under a pleiotropic model (i.e., the DNAm levels of two CpG sites are associated due to the same causal variant), we sampled a region and a causal SNP to generate DNAm level of the first CpG site using the same strategy above. The DNAm levels of the second probe was simulated based on model $\boldsymbol{m_2} = \boldsymbol{z}b_{zm2} + \boldsymbol{e}_{m2}$, where $b_{zm2}$ is the effect of the causal SNP on DNAm level of the second probe with $b_{zm2} \sim N(0, R^2_{zm2})$, $R^2_{zm2}$ being the proportion of variance in $\boldsymbol{m_2}$ explained by the causal variant, $\boldsymbol{e}_{m2} \sim N(0, \sigma^2_{e_{m1}})$ with $\sigma^2_{e_{m2}} = var(\boldsymbol{Z}b_{zm2})(1/R^2_{zm2} - 1)$. The residuals ($\boldsymbol{e}$) of the two probes were sampled from a multivariate normal distribution $\boldsymbol{e} = \begin{pmatrix} \boldsymbol{e}_{m1} \\ \boldsymbol{e}_{m2} \end{pmatrix} \sim N\left(\begin{pmatrix} 0 \\ 0 \end{pmatrix}, \begin{pmatrix} \sigma^2_{e_{m1}} & r_p\sigma_{e_{m1}}\sigma_{e_{m2}} \\ r_p\sigma_{e_{m1}}\sigma_{e_{m2}} & \sigma^2_{e_{m2}} \end{pmatrix}\right)$, where $r_p$, $R^2_{zm1}$ and $R^2_{zm2}$ were randomly sampled from the observed distributions in the LBC cohorts mentioned above. We detected the mQTL for each probe using linear regression and performed an SMR analysis to test for association between the two probes. We then performed a HEIDI test if the SMR association p-value was < 0.05, and repeated the simulation 5,000 times to evaluate the inflation/deflation of the HEIDI test-statistics under this simulation scenario (**Fig. S12d**).

**Supplementary Note 3. Acknowledgments**

**HRS (dbGaP accession: phs000428.v1.p1):** HRS is supported by the National Institute on Aging (NIA U01AG009740). The genotyping was funded separately by the National Institute on Aging (RC2 AG036495, RC4 AG039029). Genotyping was conducted by the NIH Center for Inherited Disease Research (CIDR) at Johns Hopkins University. Genotyping quality control and final preparation of the data were performed by the Genetics Coordinating Center at the University of Washington.

**UK10K (EGA accession: EGAS00001000108):** The UK10K project was funded by the Wellcome Trust award WT091310. Twins UK (TUK): TUK was funded by the Wellcome Trust and ENGAGE project grant agreement HEALTH-F4-2007–201413. The study also receives support from the Department of Health via the National Institute for Health Research (NIHR)-funded BioResource, Clinical Research Facility and Biomedical Research Centre based at Guy's and St. Thomas' NHS Foundation Trust in partnership with King's College London. Dr Spector is an NIHR senior Investigator and ERC Senior Researcher. Funding for the project was also provided by the British Heart Foundation grant PG/12/38/29615 (Dr Jamshidi). A full list of the investigators who contributed to the UK10K sequencing is available from http://www.UK10K.org.

**GTEx (dbGaP accession: phs000424.v6.p1):** The Genotype-Tissue Expression (GTEx) Project was supported by the Common Fund of the Office of the Director of the National Institutes of Health (commonfund.nih.gov/GTEx). Additional funds were provided by the NCI, NHGRI, NHLBI, NIDA, NIMH, and NINDS. Donors were enrolled at Biospecimen Source Sites funded by NCI Leidos Biomedical Research, Inc. subcontracts to the National Disease Research Interchange (10XS170), Roswell Park Cancer Institute (10XS171), and Science Care, Inc. (X10S172). The Laboratory, Data Analysis, and Coordinating Center (LDACC) was funded through a contract (HHSN268201000029C) to the The Broad Institute, Inc. Biorepository operations were funded through a Leidos Biomedical Research, Inc. subcontract to Van Andel Research Institute (10ST1035). Additional data repository and project management were provided by Leidos Biomedical Research, Inc.(HHSN261200800001E). The Brain Bank was supported supplements to University of Miami grant DA006227. Statistical Methods development grants were made to the University of Geneva (MH090941 & MH101814), the University of Chicago (MH090951, MH090937, MH101825, & MH101820), the University of North Carolina - Chapel Hill (MH090936), North Carolina State University (MH101819), Harvard University

(MH090948), Stanford University (MH101782), Washington University (MH101810), and to the University of Pennsylvania (MH101822).
